# Nature representativeness in South American protected areas: Country contrasts and conservation priorities

**DOI:** 10.1101/456558

**Authors:** Germán Baldi, Santiago A. Schauman, Marcos Texeira, Sofía Marinaro, Osvaldo A. Martin, Patricia Gandini, Esteban G. Jobbágy

**Author notes:** Corresponding author: Germán Baldi, Ejército de los Andes 950, San Luis, San Luis, D5700HHW, Argentina.

## Abstract

**Background:** South America faces strong environmental transformations due to agriculture and infrastructure expansion and due to demographic growth, demanding immediate action to preserve natural assets by means of the deployment of protected areas. Currently, 7.1% of the (sub)continent is under strict conservation categories (I to IV, IUCN), but the spatial distribution of these 1.3 x 10^6^ km^2^ is poorly understood. We evaluate protected area representativeness, map conservation priorities and assess demographic, productive or geopolitical causes of the existing protection spatial patterns using a random forest method.

**Methods:** We characterized representativeness by two dimensions: the extent and the equality of protection. The first refers to the fraction of a territory under protection, while the second refers to the spatial distribution of this protection along natural conditions. We characterized natural conditions by 113 biogeographical units (specifically, ecoregions) and a series of limited and significant climatic, topographic and edaphic traits. We analyzed representativeness every ten years since 1960 at national and continental levels. In the physical approach, histograms allowed us to map the degree of conservation priorities. Finally, we ranked the importance of different productive or geopolitical variables driving the observed distributions with a random forest technique.

**Results:** Representativeness was variable across countries in spite of its priority in conservation agendas. Brazil, Peru and Argentina underrepresented a significant fraction of their natural diversity, while Bolivia and Venezuela protected their natural diversity equitably under extensive conservation networks. As protected networks increased their extent, so did their equality across countries and within them through time. Mapping revealed as top continental priorities southern temperate, subhumid and fertile lowland environments, and other country-specific needs (e.g., hot, humid plains of Venezuela). Protection extent was generally driven by a low population density and isolation, while other variables –like distance to frontiers, were relevant only locally (e.g., in Argentina).

**Discussion:** Our description of the spatial distribution can help societies and governments to improve the allocation of conservation efforts, being top continental priorities the southern temperate, subhumid and fertile lowland environments. We identify the main limitations that future conservation efforts will face, as protection was generally driven by the opportunities provided by low population density and isolation. From a methodological perspective, the complementary physical approach reveals new properties of protection and provides tools to explore nature representativeness at different spatial, temporal and conceptual levels, complementing the traditional ones based on biodiversity or biogeographical attributes.

## INTRODUCTION

Over the last three decades, most South American countries have undergone an unprecedented expansion of cultivated lands, infrastructure and urban areas (Pacheco 2012). These changes reflect and lead to national economic growth. However, they also pose negative local to global environmental impacts, mainly associated with the loss of biodiversity and with the provision of multiple ecosystem services (Carreño et al. 2012). Under the dominant economic logic and mainly due to still great availability of resources and low population densities, this region will probably maintain or increase its role as a global supplier of raw materials (Alexandratos & Bruinsma 2012). In this sense, protected areas stand as one of the most efficient tools to protect nature in all its forms and in the long term (Hoekstra et al. 2005).

Past conservation efforts of individuals and conservation agencies have resulted in a globally significant increase in protected areas from year to year (Chape et al. 2005; Watson et al. 2014). In the early 20^th^ century, South American countries followed the seminal ideas of North America and Europe, which emphasized preserving iconic landscape features (Wirth 1962). Almost a century later, more than one thousand protected territories exist under diverse legal figures (e.g., national parks, reserves, or monuments). They encompass 7.1% of the continental surface (1.3 x 10^6^ km^2^, almost the size of Peru), a fraction slightly above the global value (Fig. 1 and Table 1). However, this growing area does not yet provide an adequate representation of natural conditions at continental or national levels (Joppa & Pfaff 2009; Baldi et al. 2017). Two factors are likely to account for this. The first is the complex interplay of motivations that lead to protection, like guarding economically valuable assets or biodiversity hotspots (McNeely et al. 1994; Watson et al. 2014). These motivations have been of variable strength through history and across territories. The second factor is associated with the limitations that different human forces (e.g., cultivation) impose over motivations, which ultimately drive protection to areas that face little human intervention and may have comparatively low opportunity-costs, at least at the time of their establishment (Joppa & Pfaff 2009; Baldi et al. 2017).

**Figure 1.**
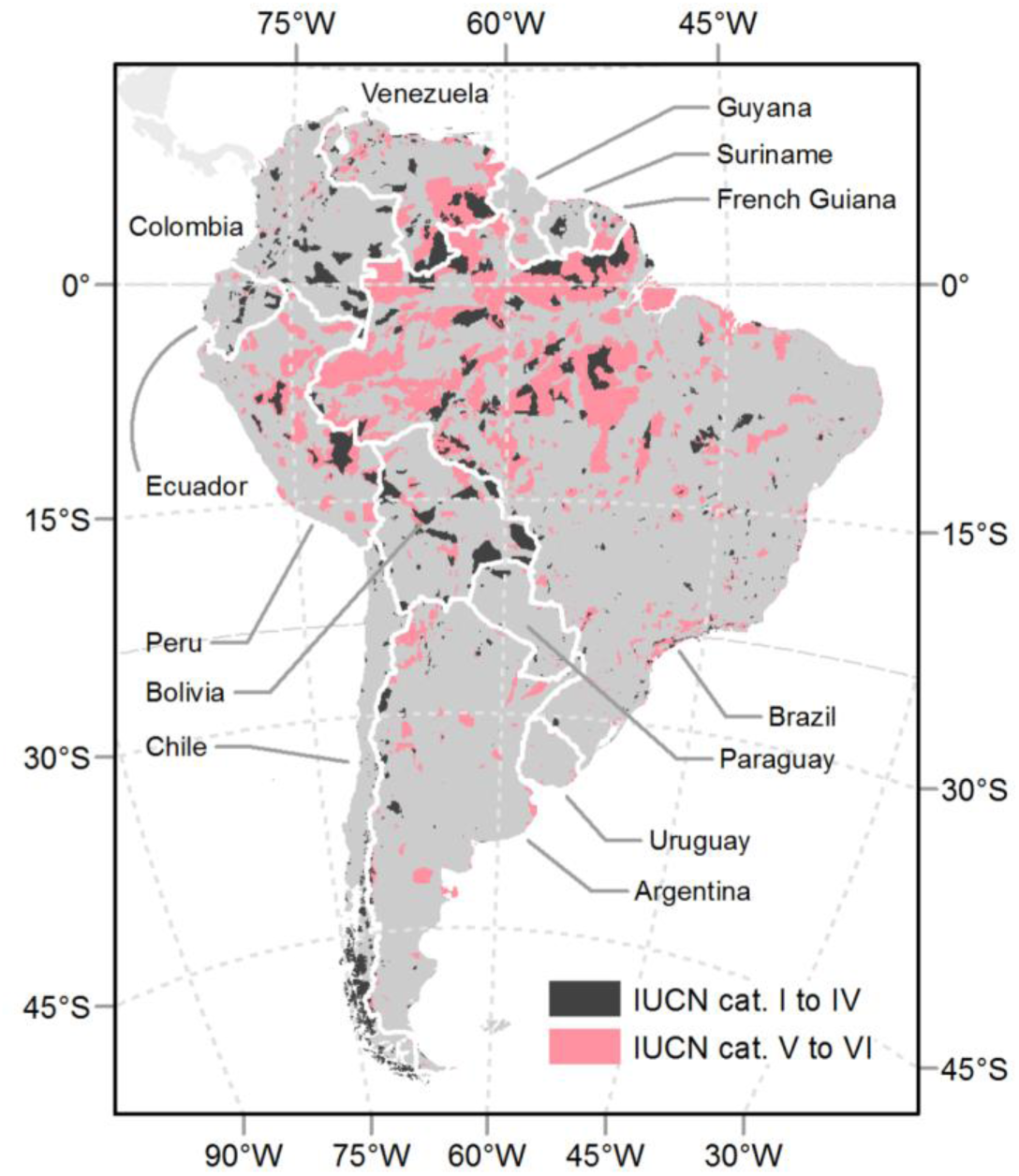
Protected areas in South America. Protected areas from the “World Database on Protected Areas” (IUCN and UNEP-WCMC 2016). Only terrestrial areas categorized as I to IV (IUCN 1994) were considered in analyses.

**Table 1.** South American networks of protected areas. General description of the networks according to the “World Database on Protected Areas”, Annual Release 2016 (IUCN and UNEP-WCMC 2016). Territories outside South America are depicted to contextualize continental values.

| **Territory** | **Area**<br>**(1,000**<br>**km<sup>2</sup>)** | **Area**<br>**under**<br>**protection,**<br>**cat. IUCN**<br>**I-IV (1,000**<br>**km<sup>2</sup>)** | **Protection**<br>**extent, cat.**<br>**IUCN I-IV**<br>**(%)** | **Number of**<br>**protection**<br>**units, cat.**<br>**IUCN I-IV** | **Area**<br>**under**<br>**protection,**<br>**cat. IUCN**<br>**I-VI (1,000**<br>**km<sup>2</sup>)** | **Protection**<br>**extent, cat.**<br>**IUCN I-VI**<br>**(%)** | **Number of**<br>**protection**<br>**units, cat.**<br>**IUCN I-VI** | **First**<br>**protected**<br>**area (yr)** |
| --- | --- | --- | --- | --- | --- | --- | --- | --- |
| Argentina | 2,793 | 64 | 2.3 | 131 | 243 | 8.8 | 360 | 1934 |
| Bolivia | 1,099 | 178 | 16.2 | 53 | 258 | 23.8 | 195 | 1939 |
| Brazil | 8,515 | 478 | 5.6 | 676 | 2,464 | 29.2 | 2,066 | 1914 |
| Chile | 756 | 139 | 18.3 | 126 | 129 | 18.4 | 149 | 1907 |
| Colombia | 1,142 | 135 | 11.8 | 86 | 156 | 13.9 | 600 | 1977 |
| Ecuador | 284 | 37 | 13 | 19 | 50 | 20 | 48 | 1959 |
| French Guiana (FG) | 91 | 4 | 4.9 | 26 | 44 | 53 | 29 | 1979 |
| Guayanas<br>(FG+GU+SU) | 470 | 20 | 4.3 | 50 | 81 | 18.6 | 83 | 1929 |
| Guyana (GU) | 215 | 0.61 | 0.28 | 3 | 19 | 8.9 | 25 | 1929 |
| Paraguay | 407 | 15 | 3.6 | 26 | 22 | 5.4 | 37 | 1906 |
| Peru | 1,285 | 85 | 6.6 | 28 | 370 | 28.8 | 212 | 1966 |
| Suriname (SU) | 164 | 15 | 9.2 | 21 | 19 | 13 | 29 | 1961 |
| Uruguay | 176 | 0.15 | 0.09 | 4 | 6 | 3.4 | 11 | 2006 |
| Venezuela | 916 | 140 | 15.2 | 45 | 383 | 42.5 | 106 | 1937 |
| All South American<br>countries | 18,321 | 1,296 | 7.1 | 1,294 | 4,244 | 23.16 | 3,950 | 1907 |
| Australia & New<br>Zealand | 7,958 | 703 | 8.8 | 14,163 |  |  |  | 1840 |
| Canada & USA | 19,379 | 1,847 | 9.5 | 9,193 |  |  |  | 1872 |
| Bhutan | 39.9 | 16.5 | 41.2 | 10 |  |  |  | 1966 |
| Costa Rica | 51.3 | 9 | 17.5 | 61 |  |  |  | 1955 |
| Kenya & Tanzania | 1,534 | 197 | 12.8 | 123 |  |  |  | 1905 |
| Scandinavia &<br>Finland | 1,261 | 113 | 9 | 6,451 |  |  |  | 1909 |
| Globe | 134,650 | 8,188 | 6.1 | 82,942 |  |  |  | 1838 |

In the same last three decades, the search for nature representativeness has been intensively promoted by conservation agencies. Representativeness implies that any network has to preserve a targeted area extent and, at the same time, has to sample all the natural conditions and of all levels of life organization with the same effort (Margules & Pressey 2000). Likewise, both the structure of this nature and its functioning may endure over time and not be altered by human interventions. In this line, the parties of the 2010 Aichi Convention on Biological Diversity (CBD) negotiated different conservation targets (SCBD 2010). Among them, the first clause of the Target 11 stipulated that at least 17 percent of terrestrial areas and inland waters needed to be included within protected networks by 2020. The second clause stipulated that networks needed to be ecologically representative, though no protection equality value was set. Although these targets do not consider local constraints and have no scientifically defined endpoints (Woodley et al. 2012), they are fundamental in stressing a global policy on how nature needs to be protected due to its intrinsic, non-utilitarian value. A rich collection of studies has evaluated the effectiveness of existing networks to represent and encompass nature considering the protection extent of biogeographical units, like ecological systems, ecoregions, biomes or realms (McNeely et al. 1994), and countries or continents (Chape et al. 2005). Juffe-Bignoli et al. (2014) stated that only 43% of the terrestrial ecoregions worldwide meet the 17% goal. In South America, though a strong diversification of protected networks has taken place since the 1960s, most biogeographical units are far below the Aichi Target 11, especially coastal areas and those with strong economic activity (Elbers 2011). Notable exceptions are the Southern Andes temperate and subpolar forests (Pliscoff & Fuentes-Castillo 2011) and the Amazonian moist forests (Jenkins & Joppa 2009).

Current conservation literature regarding representativeness reveals two theoretical and methodological biases. First, few studies address the extent of protection along continuous physical gradients like temperature o altitude (e.g., Kamei & Nakagoshi 2006; Joppa & Pfaff 2009; Baldi et al. 2017), while most describe it on the basis of predefined biogeographical units. This bias in how nature is conceived and measured is notable, given the indissoluble relationship between the physical environment and the structure and functioning of the ecosystems and given the intrinsic value of the physical environment as a constituent part of nature (Schimper et al. 1903; Holdridge 1947; Del Grosso et al. 2008; Huston 2012). Second, few studies analyze if protected areas are equally distributed within a territory, defined either by political or biogeographical units (e.g., Barr et al. 2011; Baldi et al. 2017). Given these biases, we propose a joint exploration of the two complementary dimensions of representativeness (i.e., extent and equality), considering natural conditions by means of predefined biogeographical units (a traditional biocentric approach) and physical variables (a complementary and customizable approach). We aim, thus, to provide a more comprehensive picture of any conservation status. Furthermore, the relationship between protection extent and equality is still unexplored. This information would make it possible to evaluate efficiencies and advances in representativeness under alternative settings of protected area networks.

In this paper we first characterize for South America the protection extent and equality of natural conditions within terrestrial protected areas explicitly designated for nature protection –i.e. those categorized as I-IV under the International Union for Conservation of Nature guidelines (IUCN, 1994). Second, we explore the relationship between extent and equality among countries and within them every 10 years since 1960. Third, we map conservation priorities according to the current spatial distribution of protection along physical gradients. Fourth, we relate the current spatial distribution of protection to human conditions (demographic, productive or geopolitical).

## METHODS

### Data sources

Protected areas data from South America came from the "World Database on Protected Areas", Annual Release 2016 (IUCN and UNEP-WCMC 2016). For all analyses, we considered only terrestrial areas explicitly designated for nature protection, i.e., IUCN categories I to IV (1994). In order to achieve the first three objectives, we evaluated the spatial and temporal distribution of protected areas across natural conditions, characterized by (i) 113 biogeographical units from the Olson et al. (2001) "Terrestrial ecoregions of the world" data base and (ii) the combination of five continuous physical variables representing main climatic, topographic and edaphic traits of a territory (variables #1 to #5, Table 2). For the third objective, biogeographical data was discarded from the exploration. In order to achieve the fourth objective, we evaluated the current spatial distribution of protected areas across human conditions related to different motivations of conservation (McNeely et al. 1994; Watson et al. 2014) (variables #6 to #10,Table 2). Specifically, "tourism attractiveness" quantifies the influence of aesthetic/recreational values of a territory in the implementation of protected areas. The "distance to frontiers" depicts the importance of protection close to international borders, conceiving these as territories where is necessary to assert sovereignty in a peaceful manner. The last three variables, "population", "distance to roads" and "cropland suitability", quantifies the protection in territories that have a low economic value for traditional and profitable land uses (Baldi et al. 2017).

**Table 2.**
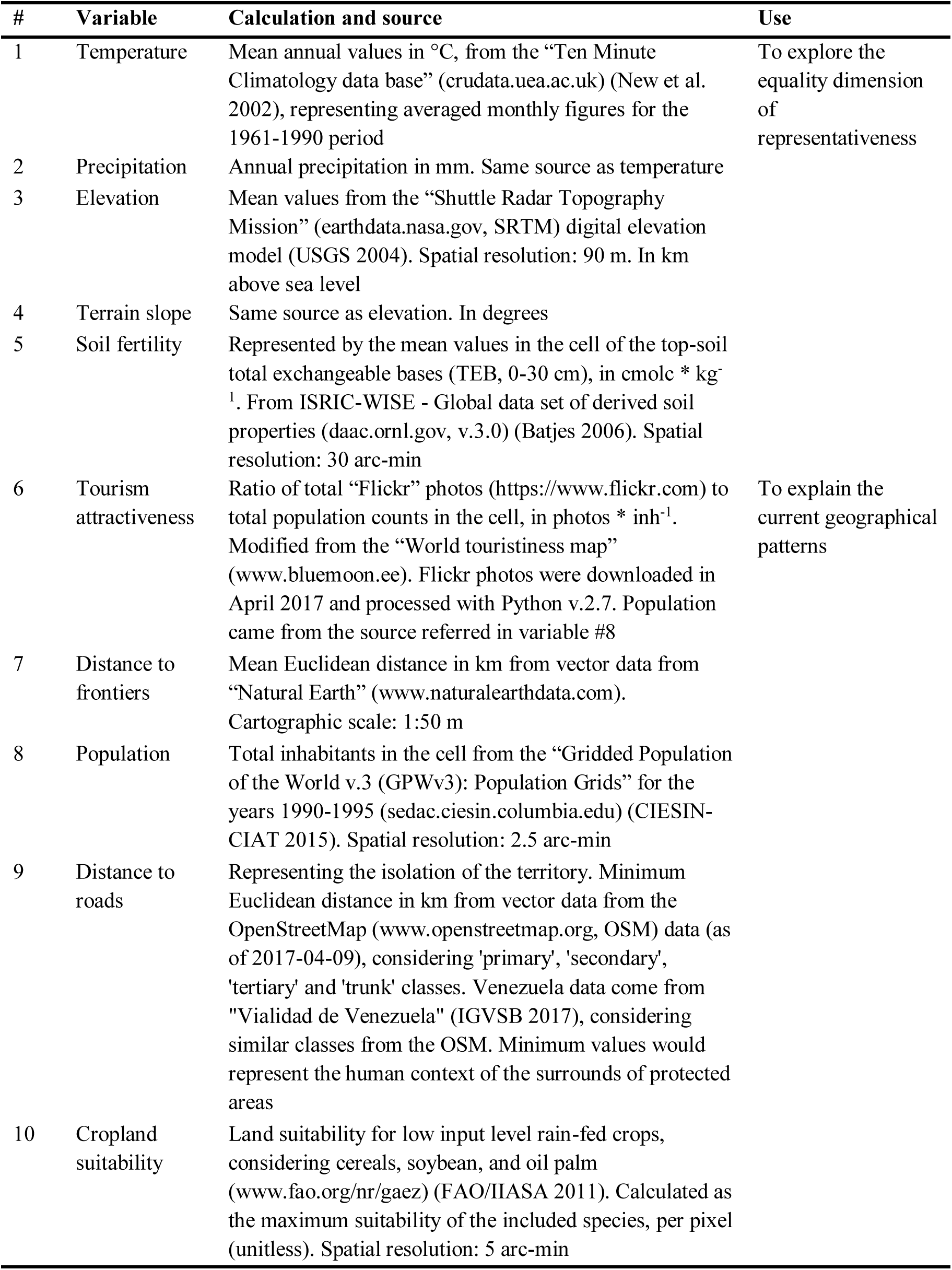
List of physical and human variables. List of physical (i.e., climatic, topographic or edaphic) and human (i.e., demographic, productive or geopolitical) variables used to evaluate the distribution of protected areas.

### Sampling procedure

We explored the spatial and temporal distribution of protected areas at national and continental levels (Fig. 1). The Guayanas, i.e. French Guiana, Guyana and Suriname, were treated as a single unit due to their relatively small size and physical homogeneity.

We sampled data following two complementary approaches. In the first, protection, physical and human data were summarized into a grid of 55,414 cells of 0.1° x 0.1°. In the second, only protection data was summarized into the above mentioned biogeographical units. The first approach allowed us to analyze representativeness and map conservation priorities upon a physical basis (first three objectives) and to assess human drivers of conservation (fourth objective). The second approach allowed us to analyze both dimensions of representativeness upon a biogeographical basis (first two objectives).

Compared to other sampling approaches in which each unit of protected area is treated as a single sample, gridding offers the advantages of (i) providing a unified spatial resolution for all variables, (ii) encompassing the full range of physical and human conditions, (iii) avoiding the averaging of these conditions within very large protected areas and (iv) providing a representation of the context of protected areas by characterizing the full grid cell in which they are embedded (97.5% of the cells incorporate unprotected conditions).

## Data analysis

For the first sampling approach we generated 840 histograms –i.e., 12 territories * 10 continuous variables * 7 temporal periods–, containing three sets of information: (i) the absolute extent under protection (in 1,000 km^2^) of each *j* class (interval in the histograms) of the *i* continuous variable, (ii) the relative extent under protection (in %) of the class *j* of the *i* continuous variable –PEx_ij_ and (iii) the extent (in 1,000 km^2^) of each class of the *i* continuous variable (Fig. 2). In those cells shared by two or more countries, PEx_ij_ values corresponded exclusively to the focus country. All histograms had 10 bins or classes, regardless the variability of the *i* continuous variable in the territory (i.e., a country or the entire continent). In order to avoid long tails in the histograms, lower and upper *j* classes were grouped using the percentile values 0.025 and 0.975 of the *i* continuous variable. Exclusively to assess the stability of the standardized binning method, we also calculated G' considering a variable number of classes following the Sturges approach (1926). We conducted the statistical analyses corresponding to the first three objectives with the PEx_ij_ values; the two remaining sets of information were shown only for descriptive purposes.

**Figure 2.**
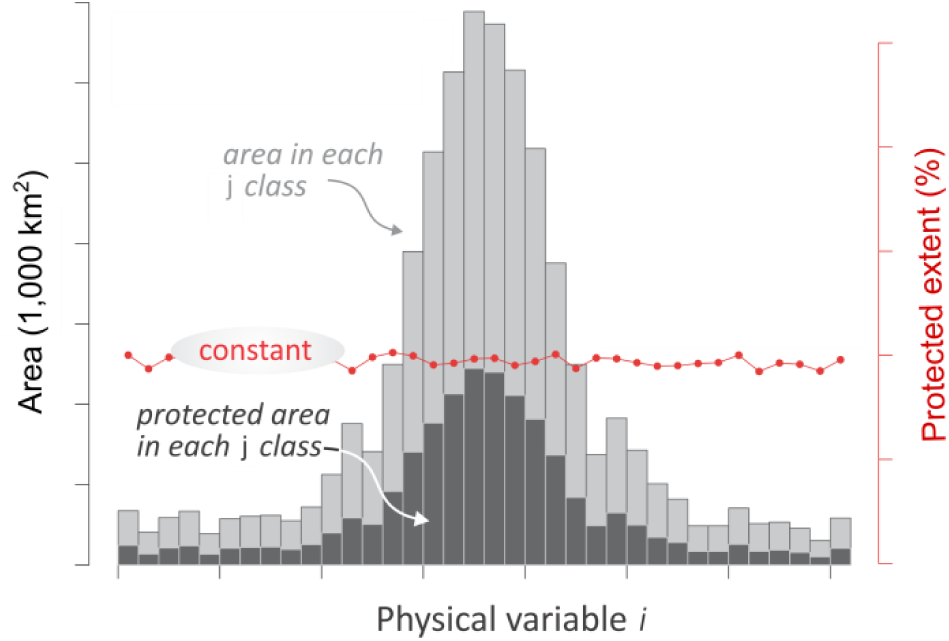
Expected protection pattern according to the representativeness motivation. Model of a completely balanced protected area network along a *i* physical continuous gradient. The encircled text refers to the expected and tested behavior. Three measurements are shown in the histogram: the extent in each class –intervals– of the continuous variables (light gray bars), the absolute extent under protection in each class (dark gray bars), and the relative extent under protection in each class (red dots –PEx_ij_, and lines).

In order to explore protection equality, we analyzed for each territory the spatial and temporal distribution of PEx_ij_ values along the five *i* continuous physical variables (Table 2) by means of the Gini coefficient (G_i_) (Barr et al. 2011; Chauvenet et al. 2017). As G_i_ measures the inequality among values of a frequency distribution, we calculated its reverse (1 - G; hereafter, G_i_'). If all PEx_ij_ values are equal for all *j* classes of the *i* continuous physical variable, G_i_' achieves a maximum equality of value 1, independently of the PEx_ij_. If the difference in PEx_ij_ values increase, G_i_' decreases to a theoretical minimum of 0.

At last, we achieved a single G' value by territory by averaging the five G_i_' values. However, in this averaging we reduced the effects of multicollinearity between continuous physical variables by eliminating from the analysis an *i* variable if the module of its correlation coefficient with the *i+1* variable resulted greater than 0.55, according to a Kendall’s τ non-parametric test (Whittaker 1987) (Fig. S1). For the second sampling approach (biogeographical) a G' was also calculated, but PEx values were calculated on each predefined biogeographical unit within each territory.

In order to map the priorities of conservation (Pr, in %) exclusively based on the *i* continuous physical variables, we analyzed the difference (in percentage) to a condition of absolute protection (100% of the area) following the equation:

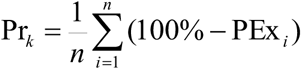

where *k* represents the grid cell in each territory and *n* the number of continuous physical variables. Again, we removed *i* variables with a |τ| > 0.55. In this sense, those grid cells that have low PEx_i_ values will have a large difference to a 100% protection condition and therefore, will achieve a higher Pr.

Finally, histograms allowed us to characterize the behavior of the protection extent along human gradients (variables #6 to #10, Table 2) in terms of shape and sign, but not to rank the relative importance of these variables as causal drivers. In this sense, for the fourth objective, we applied exclusively for the current protected area data of each individual cell (up to 2016), a random forest algorithm (Breiman 2001) which estimates the variable importance (VI) by looking at how much the mean square error (MSE) increases when the out-of-bag data (OOB) for that variable is permuted while all others are left unchanged (Liaw & Wiener 2002). This technique has been extensively used to solve problems of classification and regression in ecology and related disciplines (Cutler et al. 2007). The allocated VI can differ substantially with the selection of number of trees to grow (*ntree*), the minimum size of the terminal nodes (*nodesize*), or the number of input variables at each split (*mtry*) (Grömping 2009). We chose those values that minimize the OOB-MSE of the model (*ntree* = 500 and *nodesize* = 1). The VI was used here with an explanatory and interpretative rather than predictive aim. All calculations were run in RStudio v. 1.0.143 (RStudio Team 2018) (packages foreign, ggplot2, ggrepel, gridExtra, ineq, lattice, png, Segmented) and Python v.2.7 (packages Scikit-learn, Numpy) (Pedregosa et al. 2011; van der Walt et al. 2011).

## RESULTS

Beyond the more evident differences in protection extent, with Chile, Bolivia and Venezuela at the top of the ranking (Table 1), South American countries strongly contrasted in the equality of protection of their natural conditions (Fig. 3). The Guayanas, Bolivia and Venezuela achieved the highest equality values when natural conditions were represented by biogeographical units (G' > 0.40). The variability reached 1.9 times between the leading country (the Guayanas) and the last country in the ranking (Argentina) (Fig. 3a). However, Bolivia, Colombia, Venezuela and Ecuador achieved the top of the ranking when natural conditions were represented by physical variables (G' > 0.72) (Fig. 3b). Bolivia and Colombia showed an equality of physical conditions 3.5 times higher than Uruguay and 1.5 times higher than Argentina (the second country in a bottom-up ranking).

**Figure 3.**
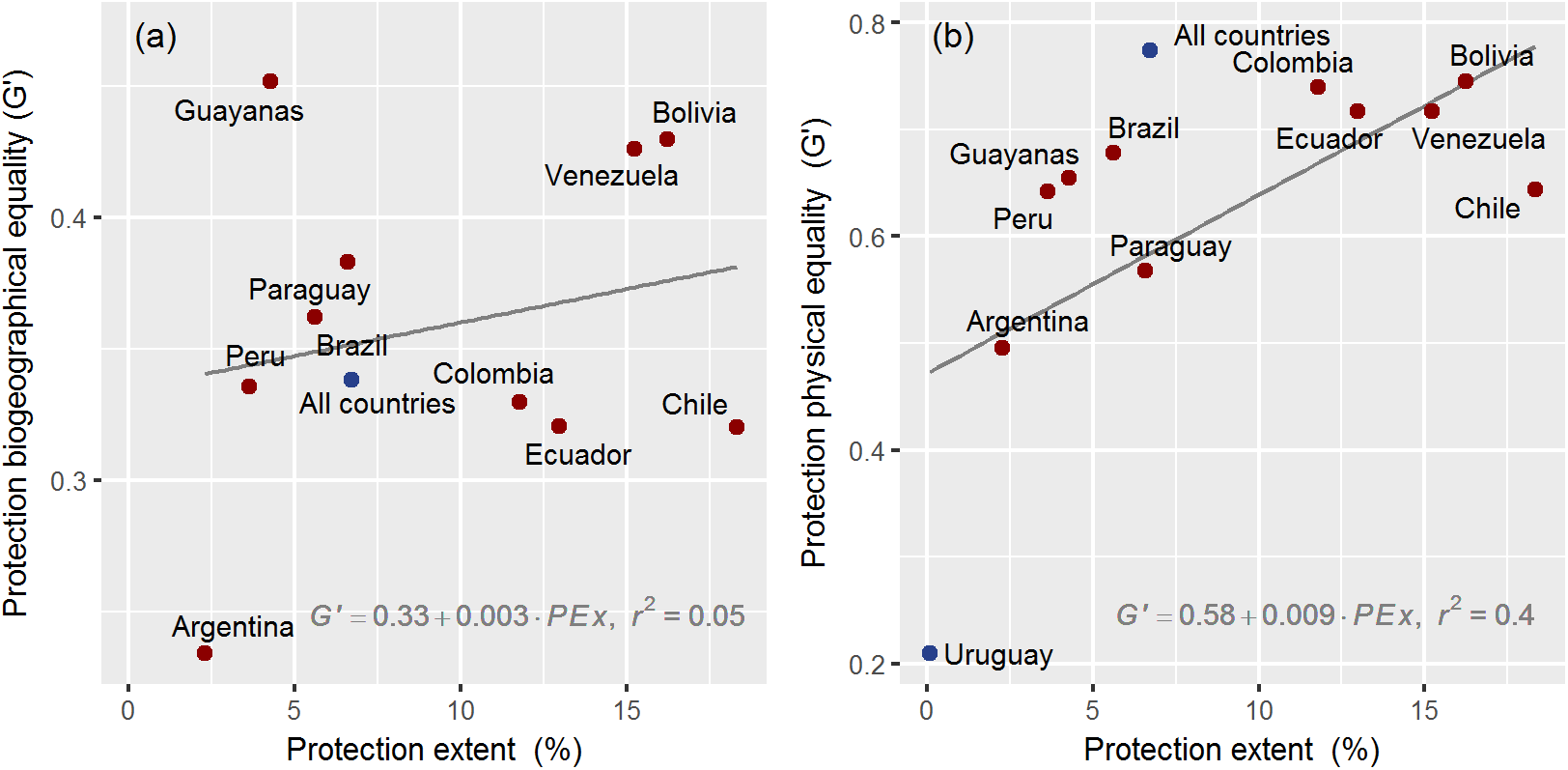
Current representativeness. Relationship between current protection extent (%) and equality (G') in South America. In (a) equality is calculated on the basis of biogeographical units and in (b) on the basis of physical continuous variables. Continental and Uruguayan results (in blue) did not feed linear regressions. As Uruguay has only one ecoregion, no equality value is quantified in (a) panel.

When considering biogeographical units, we found no relationship between the two analyzed dimensions of protection (G' = 0.33 + 0.003 · PEx, r^2^ = 0.05). As examples, the Guayanas and Argentina reached contrasting equality values despite both territories having a low and relatively similar protected extent (4.3% and 2.3%, Table 1), while Chile and Peru reached a similar equality (G' ≈ 0.32) with a different protected extent. Instead, when considering physical variables, we found a weak relationship in terms of slope, but a strong one in terms of coefficient of determination (G' = 0.58 + 0.09 · PEx, r^2^ = 0.4). Notably, Chile, in spite of having the highest protection extent among countries (18.3%, Table 1), attained from medium to low equality (G' = 0.64). No notable changes occurred in the above mentioned patterns when a different binning method was applied in the generation of histograms (Fig. S2).

The above-described patterns were unrelated to the date in which each country created its first protected area, as shown by the new and old networks of Colombia and Venezuela (1977 vs. 1937, Table 1). Historically, as the extent of protected networks increased so did their equality (Fig. 4). However, we did not identify a typical or synchronized trend of representativeness in the last 5 decades, even though most countries reached their maximum equality between 1990 and 2000. The individual behavior of countries was characterized either by a general parallel increase in both extent and equality (i.e., small in Argentina, or large in Ecuador), a steady increase in extent but not in equality (i.e., in Chile), or an increase in the extent which eventually reduces equality. Additionally, decade-to-decade changes in both dimensions of representativeness did not follow a pattern of acceleration or stabilization. We found that the rates of increase in equality generally diminished in comparison with the rate of creation of new protected areas, with notable exceptions in Argentina and Paraguay during the 1980-2010 period (Fig. S3). The biogeographical approach to calculating equality offered a similar historical pattern, but with large differences among countries (Fig. S4).

**Figure 4.**
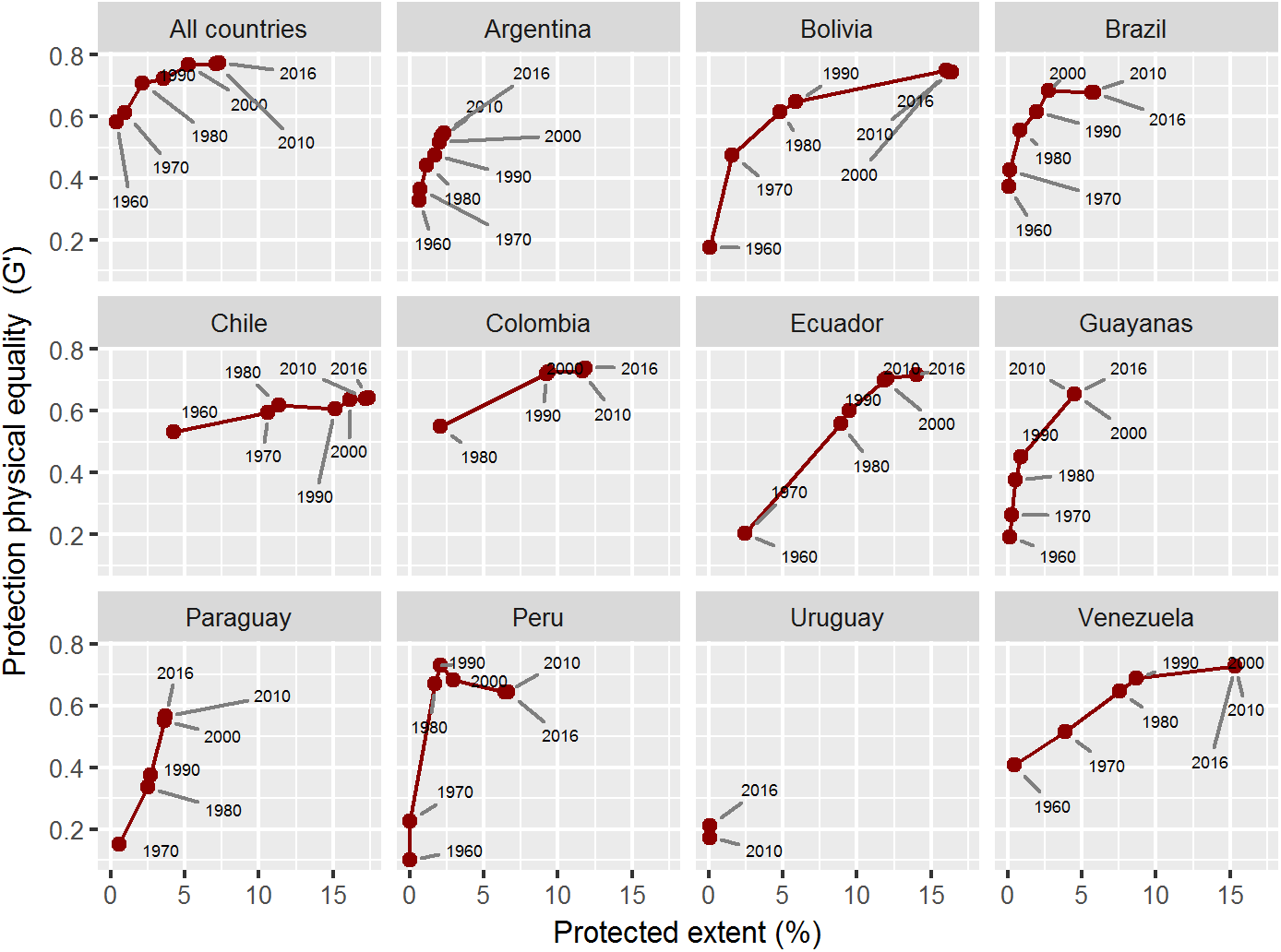
Evolution of representativeness based on physical variables. Temporal evolution of the relationship between protection extent (%) and equality (G') in South America. Equality is calculated on the basis of physical continuous variables. Each dot indicates the end point of a temporal period except for the 1960 one, which indicates the data before 1960, inclusively.

Based on the current spatial distribution of protection along physical gradients, maps revealed continuities and discontinuities of conservation priority among countries (Figs. 5a and S5 and S6). When priority areas match ecoregions, we mention them due to their popularity in the scientific literature. The hot, humid plains of the Llanos and the surrounding broadleaf forests in Colombia and Venezuela are an example of a transboundary natural system that deserves more attention. The Ecuadorian and Peruvian Andes and the subhumid highlands of the Atlantic forest in Argentina, Brazil and Paraguay, are examples of divergent conservation needs. Generally, most national priorities coincide with temperate subhumid areas originally covered by grasslands and hot arid to semiarid areas originally covered by deserts, shrublands, savannas and dry forests. A detailed description of the national priorities is depicted in Table S1. Continental priorities were the subtropical and temperate plains and plateaus of Argentina and Uruguay; and the coastal dry areas of the Pacific coast, northeastern Brazil and northwestern Venezuela (Figs. 5b and S5). The South Andes and the humid tropical lowlands of the Amazonian basin achieved significant better protection levels, with numerous, very large protected areas (e.g. Jaú National Park in Brazil and Bernardo O'Higgins National Park in Chile).

**Figure 5.**
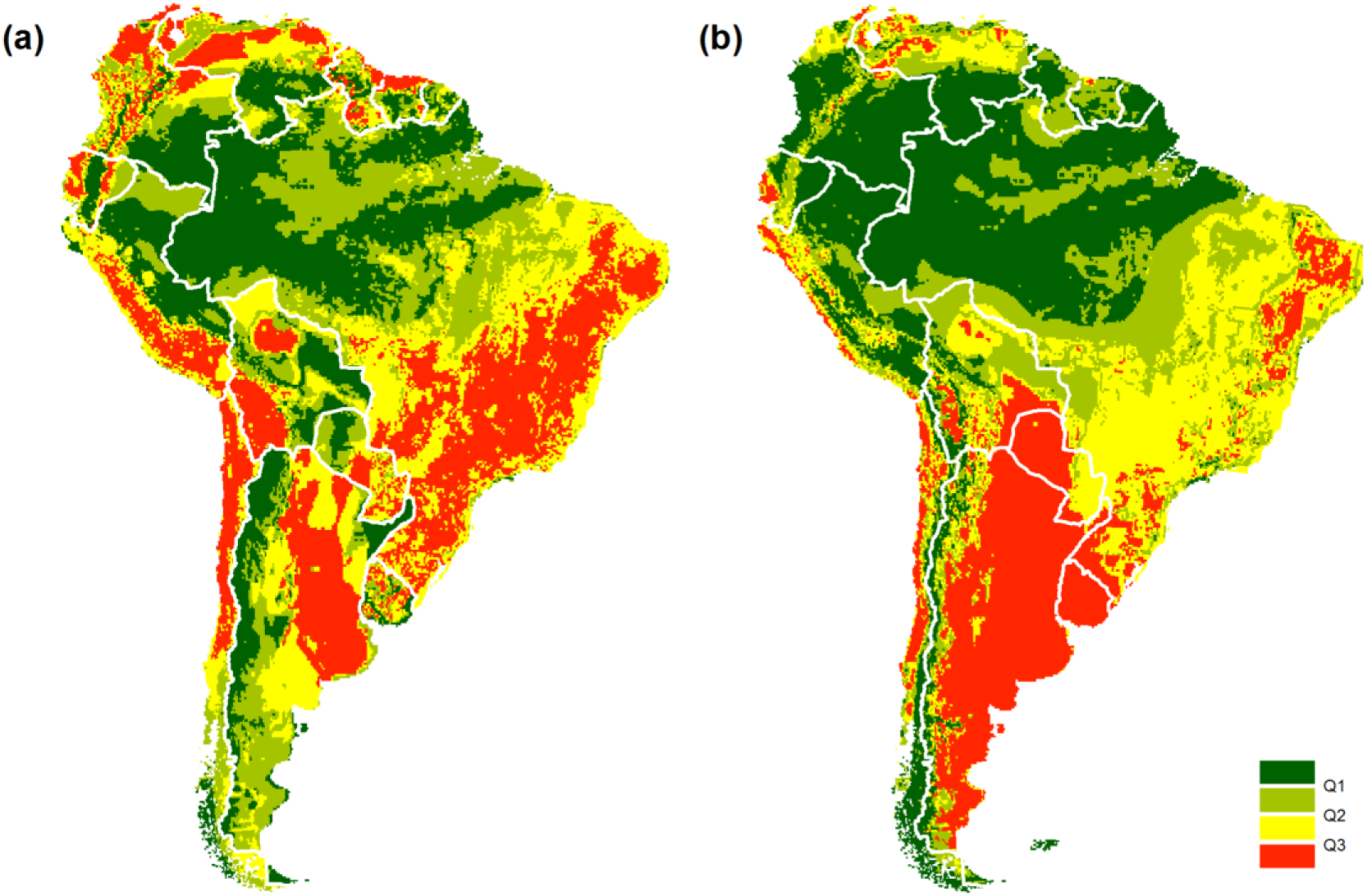
Conservation priorities. (a) National conservation priorities (Pr) in South America according to the current spatial distribution of protection extent along physical gradients. (b) Same as previous, but considering the continent as a single unit. Pr data is classified into quartiles (Q), i.e. each class encompasses a quarter of the grid cells. Red represents the highest priority, dark green the lowest. White lines represent national divisions. Detailed maps are presented in Fig. S5.

The current spatial distribution of protected areas was related to the historical limitations to conservation that demographic, productive, or geopolitical forces imposed (Figs. 6, S5 and S6). According to the random forest analysis, the most important human variable explaining the spatial distribution of protected areas was population, followed by distance to roads, distance to frontiers, cropland suitability and tourism attractiveness (from VI = 27.1 to VI = 10.7, by averaging countries; Table 3). Protected areas were preferentially allocated in sparsely populated areas, especially in Peru, where its importance was 20% higher than in Paraguay (the next country on the list). The farther a territory was from a road, the greater the protection level. In Chile, the distance to roads achieved a maximum relevance, with a magnitude 1.5 times higher than in Colombia, the next country in the ranking (VI = 39 and 26.9, respectively). Protected areas were comparatively closer to international frontiers, with significant strength in Argentina and Brazil (VI = 24.9 and 24.6, respectively). The effect of cropland suitability on protection was dissimilar among countries (positive or negative relationships). Remarkably, in Argentina and Chile, cropland suitability had a strong negative relationship with the spatial distribution of protected areas, but a relatively small importance according to the random forest. Finally, tourism attractiveness seems to have played a role driving conservation in Uruguay (VI = 31.2) and in Argentina, Chile and Ecuador (VI ≈ 14).

**Figure 6.**
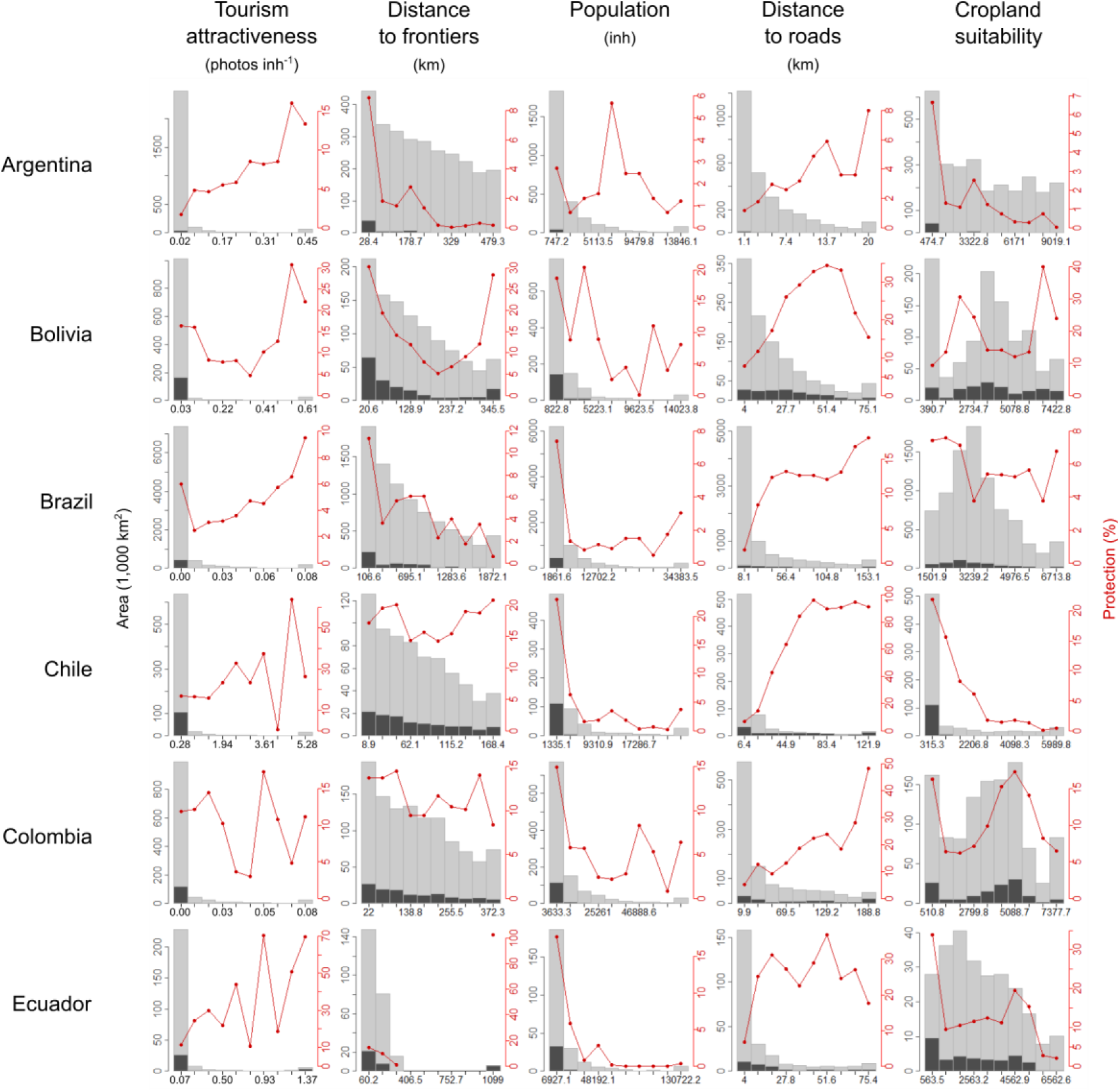
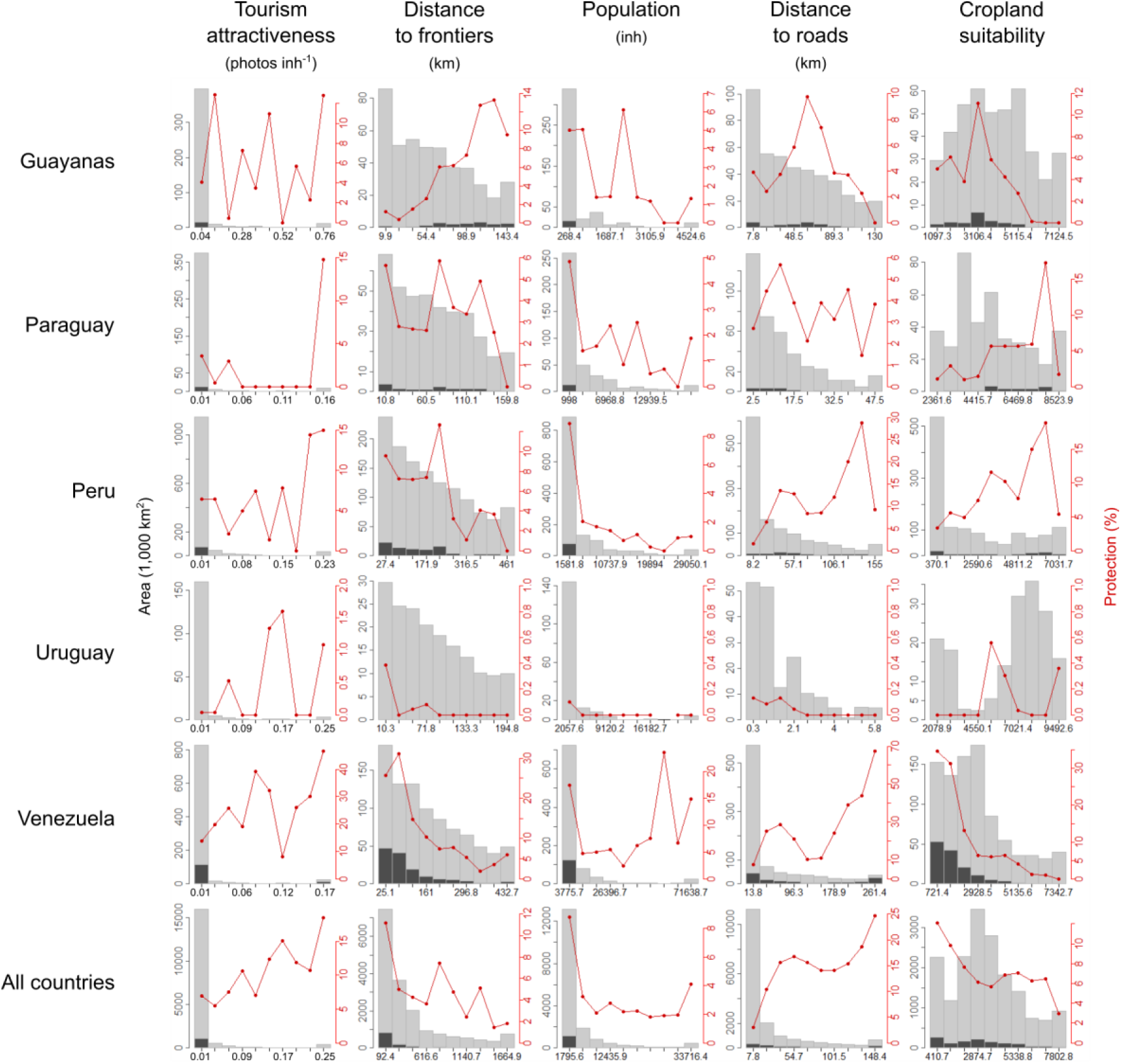
Drivers of protected areas. Current spatial distribution of protection extent along human gradients (demographic, productive, or geopolitical) in South America. See graphic explanations in Fig. 2. Lower and upper *j* classes were grouped using the percentile values 0.025 and 0.975 of the *i* continuous variable.

**Table 3.** Variable importance according to a random forest. Legend: Relative importance (% MSE) of five variables by territory according to the random forest analysis.

| **Territory** | **Tourism**<br>**attractiveness** | **Distance to**<br>**frontiers** | **Population** | **Distance to**<br>**roads** | **Cropland**<br>**suitability** |
| --- | --- | --- | --- | --- | --- |
| Argentina | 14.6 | 24.9 | 27.8 | 18.1 | 14.6 |
| Bolivia | 6.3 | 22 | 29 | 20.9 | 21.9 |
| Brazil | 3.9 | 24.6 | 29 | 21.9 | 20.5 |
| Chile | 13.4 | 15.2 | 25.3 | 39 | 7.1 |
| Colombia | 5.7 | 21 | 24.9 | 26.9 | 21.5 |
| Ecuador | 14.1 | 20.7 | 23.6 | 19.6 | 22.1 |
| Guayanas | 8.4 | 22 | 27.5 | 20.8 | 21.3 |
| Peru | 5.9 | 18.3 | 35.9 | 18.1 | 21.9 |
| Paraguay | 6.2 | 19.4 | 29.9 | 22.7 | 21.8 |
| Uruguay | 31.2 | 20.2 | 19.2 | 6.2 | 23.2 |
| Venezuela | 8.4 | 18.9 | 25.8 | 22.8 | 24.1 |
| Country average | 10.7 | 20.7 | 27.1 | 21.5 | 20.0 |
| All South<br>American<br>countries | 6.1 | 21.6 | 27.8 | 24.2 | 20.4 |

## DISCUSSION

Representative coverage of nature by protected areas is far from being achieved in most of South America considering areas explicitly designated for protection (i.e., IUCN I to IV –1994), in spite of being an explicit priority in national and international agendas (e.g., SCBD 2010). Extensive and physically and ecologically diverse countries like Brazil, Peru or Argentina, accounting for almost 70% of the continental area, still underrepresent a significant fraction of their nature (G' < 0.68 when considering physical variables). Colombia, Bolivia, Venezuela and Ecuador perform better, surpassing the 12% of protection and converging in the equality of natural conditions (G' ≈ 0.73 by applying the same approach). According to Barr et al.’s ranking (2011), Bolivia, Venezuela and Ecuador achieved an equality of protection 25% lower than Costa Rica, or 50% lower than Bhutan (countries renowned for their pioneering conservation efforts). As a whole, South America slightly surpasses the global protection extent (7.1% vs. 6.1%, Table 1), but is still 10% under the Aichi Target 11. Regarding protection equality, the continent occupies a low to intermediate position, below Europe, Central Asia and Sub-Saharan Africa (Baldi et al. 2017).

Our results suggest that the chance of balancing natural conditions increases by expanding the extent of protected networks. This positive relationship arises both in the spatial comparison between countries (Fig. 3b) and in the temporal comparison within countries (Fig. 4). Models suggest a general stabilization or even a decrease in the equality value as networks expand. A tentative explanation for this is that as protected networks expand, underrepresented conditions become scarcer (Marinaro et al. 2012). On the one hand, Chile illustrates the stabilization behavior, probably due to the difficulties to balance a network in which large tracts of subpolar forests were initially incorporated (near half of today's Magellanic ecoregion is under public protection). On the other hand, the historical trends of representativeness in Peru and Brazil exemplify the equality decline behavior, as a result of the incorporation of large or numerous protected areas over already well-represented conditions. Together, these two countries hold almost three-quarters of the iconic Amazon rainforest, a natural system that has attracted strong protection efforts in the second half of the 20^th^ century (Peres & Terborgh 1995). According to Jenkins and Joppa (2009), this region accounted for most of the global protected area expansion of the 2000 decade. Additionally, our results for South America question Barr et al.’s (2011) suggestion about a general inverse relationship between extent and equality. Beyond the different geographical space encompassed, discrepancies may be related to the biogeographical approach that they followed, as we found no association between both representativeness dimensions using this sampling approach (Fig. 3a).

Multiple reasons may explain why countries that share strong cultural and historical traits perceive and effectively protect their nature in such different ways (Table 1 and Fig. 3). Recently, we suggested that geographical differences in the distribution of protected areas were likely to reflect the interactions among policies and economy (e.g., economic context), social organization (e.g., ONG and philanthropic actions) and moral considerations (e.g., religion) (Baldi et al. 2017). In this sense, some narratives and quantitative studies describing the roots of South American protected areas highlight the effects of the (asynchronous) occurrence of financial surpluses and the alternation between government types (Barker 1980; Pauchard & Villarroel 2002; Marinaro et al. 2012; Leal 2017). According to them, autocratic governments and young democracies previous to 1970 prioritized aesthetic/recreational values, geopolitical hotspots or potential forest production, while from the 1970s onwards, the effective protection of emerging representativeness and biological conservation values was tied to financial surpluses. We showed that early and modern protected areas were established preferentially in isolated or sparsely populated territories with reduced agricultural capacity and close to international borders (Table 3 and Fig. 6) (Baldi et al. 2017). In this regard, setting aside new land to preserve biodiversity is likely to succeed in areas of comparatively low opportunity costs from economic activities, but certainly representativeness collides with agriculture, forestry and mining.

If intentions to enhance the representation of natural conditions had prevailed and had received adequate financial support in the last four of five decades, national administrations would not have been able to reverse the strong initial bias in the spatial distribution of protected areas (Figs. 4 and S4). Modern protected areas tend to be of small size and follow isolated rather than systematic conservation efforts (Marinaro et al. 2012). We consider that historical conservation patterns could be attributed to national rather than to international policies. This may be accounted for by the temporal trends in representativeness, unique for each country. Radeloff et al. (2013) support this observation by showing that advances in the protection extent were unsynchronized among countries, occurring during brief periods named "hot moments" (a concept extensible to equality). Some of these moments occurred in the first years of national networks, as in the case of Argentina in the thirties/fifties (Marinaro et al. 2012), Chile in the sixties (Pauchard & Villarroel 2002), or Colombia in the seventies (Leal 2017). We found that from 2010 (Conference of the CBD Parties) to 2016 (last year in our assessment), only 32,000 km^2^ of new protected areas were added to the continental network under categories I to IV. Therefore, with less than three years until the 2020 Aichi Target ought to be accomplished, we offer maps that help conservation agencies focus their efforts while following closely the criterion of representativeness (Figs. 5 and S5). The temperate to subtropical shrubby-to-grassy plains of eastern Argentina and Uruguay (coincident with the Pampas and Chaco ecoregions) account for the largest territory of high continental and national priority. It is worth mentioning that even though the Pampas and Chaco have diverged in their land use history, they converge in threats and challenges to conservation (Brown et al. 2006; Overbeck et al. 2015). The relatively low protection of the natural conditions hosted by these two countries also results from the fact that low- rather than high-latitude conditions prevail in South America. Near the tropics, national priorities coincide with dry systems, which have recently caught the attention of scientists and conservationists due to their unappreciated biodiversity, rapid changes and high agricultural potential (Grau et al. 2014). In contrast, disagreements between continental and national maps reveal that priorities in one country might entail oversized efforts if these conditions are uncommon in it, but well represented in neighboring countries. Likewise, well represented national conditions can still deserve further conservation efforts if those are underrepresented in neighboring countries. This can be the case of the well-protected subhumid highlands in Argentina vs. the situation in Brazil and Paraguay (Atlantic forest ecoregion) (Huang et al. 2009; Henriques 2011).

Spatially-explicit priorities like the ones we mapped can help conservation agencies focus future efforts. However, priority maps do not consider real world limitations, namely: the persistence of natural vegetation remnants, their structural and functional condition and their level of isolation, land tenure or acquisition costs. Natural vegetation has been totally removed or profoundly transformed in large tracts of the continent, with urban settlements and croplands occupying 22% of its area (Ellis & Ramankutty 2008). These same restrictions must have faced the promoters of the first protected areas. As an example, the intense use of the territory in the Argentinean and Uruguayan Pampas (Colomé 2009) or the Brazilian Atlantic forests (Henriques 2011) precedes the great transformations that took place in the last decades.

We should acknowledge that the role of additional protected areas not considered in our analysis may greatly alter results (Fig. 1 and Table 1). Previous assessments for Argentina (APN 2007), Bolivia (Larrea-Alcázar et al. 2016), Brazil (Rylands & Brandon 2005), Chile (Pliscoff & Fuentes-Castillo 2011), Guyana (Bicknell et al. 2017) and Peru (Shanee et al. 2017) highlight the importance of these areas in the achievement of targets and agreements. Aichi Target 11 did not explicitly specify under which categories this specific area should be encompassed (Woodley et al. 2012). Although IUCN categories V and VI have a dual role in promoting the preservation of biodiversity and the local economic welfare, a statutory limit for resource exploitation is not stipulated (Shafer 2015). Besides, these governmental, communal or private protected areas potentially lack formal protection and management and have uncertain conservation objectives and long-term capabilities (Shafer 2015). The effect of considering all categories becomes critical in the so-called "developing" countries, as predictions state that over the next decades, new areas will be designated under multiple-use rather than under strict categories (McDonald & Boucher 2011; Shafer 2015).

Using biogeographical units as a basis to calculate representativeness has served to report conservation progress and to derive international policies (McNeely et al. 1994). However, the proposed (gridded) physical approach reveals new properties of protection and provides tools to explore nature representativeness at different spatial, temporal and conceptual levels. First, it is sensitive to the estimation of the progress of conservation, since equality and extent are associated (Fig. 3b). Second, it allows mapping priorities at spatial resolutions which are only constrained by the available physical data. Third, the physical approach allows customizing the characterization scheme of "natural conditions" to any specific need. The selection of physical variables can be modified, expanded or improved with new or more suitable options. Fourth, it considers shifting physical patterns resulting from natural- or human-induced causes. Thus, geographical priorities could be forecasted under different climate scenarios (Scott et al. 2002; Theobald et al. 2015). Fifth, it avoids the over-inflation of the equality metric when the number of classes is low (Chauvenet et al. 2017), as the number of physical intervals is user-defined. At last, there are theoretical reasons and empirical evidence that show that physical variables are efficient estimators of the spatial distribution of species (Margules & Pressey 2000). Thus, our physical approach undoubtedly complements the traditional ones based on biogeographical attributes as well as those ones dealing with the gaps in biodiversity, from genes to ecosystems (Rodrigues et al. 2004).

## CONCLUSIONS

In this paper, we found that even the top national conservation networks are far from being saturated and balanced, as protected areas tend to be located in sparsely populated and isolated territories. We also found that by expanding the extent of protected networks, the representation of natural conditions generally increases. However, temporal trends in conservation showed that equality is not gaining strength, contrary to the dominant conservation logic. Spatially-explicit priorities can guide agencies to focus their efforts, but demographic and productive limitations are imposed on the deployment of new areas. In this sense, representativeness will only be strengthened if coupled with economic interests based on the provision of goods and services (including tourism).

## ACKNOWLEDGEMENTS

We would like to thank H.R. Grau, S. Aguiar and M. Burgman for their ideas and collaboration in different stages of the study. We also appreciate the English assistance provided by the Gabinete de Asesoramiento en Escritura Científica en Inglés, UNSL.

